# Niche partitioning in a cyanobacterium through divergence of its novel chlorophyll *d*-based light-harvesting system

**DOI:** 10.1101/2024.01.17.576076

**Authors:** Nikea J. Ulrich, Gaozhong Shen, Donald A. Bryant, Scott R. Miller

## Abstract

The evolution of novel traits can have important consequences for biological diversification. New ecological opportunities provided by a novel trait can trigger subsequent trait modification or niche partitioning; however, the underlying mechanisms of novel trait diversification are still poorly understood. Here, we report that the innovation of a new chlorophyll (Chl) pigment, Chl *d,* by the cyanobacterium *Acaryochloris marina* was followed by the functional divergence of its light-harvesting complex. We identified three major photosynthetic spectral types based on Chl fluorescence properties for *A. marina* laboratory strains, with shorter and longer wavelength types more recently derived from an ancestral intermediate phenotype. Members of the different spectral types exhibited extensive variation in the Chl-binding proteins as well as the Chl energy levels of their photosynthetic complexes. This spectral type divergence is associated with differences in the wavelength dependence of both growth rate and photosynthetic oxygen evolution. We conclude that the divergence of the light-harvesting apparatus has consequently impacted *A. marina* ecological diversification through specialization on different far-red photons for photosynthesis.

## Introduction

Novel traits are engines of biological diversification. Novelties such as new structures are associated with changes in both genotype and phenotype that often lead to changes in ecological function (1, 2). The evolution of a novel trait can enable an organism to access unoccupied niches where there are new resources to exploit (3–5), which can trigger an adaptive radiation through subsequent trait modification (6). For example, the innovation of the beetle horn as a secondary sexual trait prompted an impressive radiation of diverse horn size, shape, and number, both within and between species (7). Similarly, the evolution of eyespots on butterfly wings, which provided innovations in mate recognition and predator defense, set the stage for the dramatic diversification of wing patterns in Lepidoptera (8–10). During the process of novel trait diversification, populations might begin to partition the new niche space to avoid intraspecific competition (11). This partitioning may be temporal (12, 13), spatial (14), or with respect to resource utilization (11). The latter involves character displacement, i.e., an evolutionary change in a morphological, physiological or other trait that enables specialization on a specific resource (15). However, despite their importance, with few exceptions (e.g., 16, 17), we rarely know the underlying mechanisms of novel trait diversification.

One of the most consequential innovations in the history of life was the invention of Chl *a*-based photosynthesis by cyanobacteria, which sparked the oxygenation of Earth. Since the origin of Chl *a* and its subsequent acquisition by diverse eukaryotes via endosymbiosis, different organisms have extended the reach of the photosynthetic apparatus through the diversification of light-harvesting complexes (LHC) that harvest wavelengths of light that are inaccessible to Chl *a*. While photosynthetic reaction center structure, composition and function have been highly conserved during billions of years of Chl *a*-based oxygenic photosynthesis (18), spectrally distinct LHCs have independently evolved in diverse photosynthetic lineages over the same period of time. These independently derived LHC include the phycobilisomes (PBS) of most cyanobacteria and red algae (19, 20), the prochlorophyte Chl *a*/*b* binding proteins (Pcb) (21), the LHC proteins of green algae and plants (22, 23), and the fucoxanthin Chl *a*/*c*-binding proteins of stramenopiles and haptophytes (24, 25). LHC are often tuned to specialize on the prevailing light available in the environment (e.g., the diversity of PBS pigments in marine *Synechococcus* at varying depths; 26). LHC systems can also differ dramatically in the wavelengths they harvest. For example, PBS absorb light ranging from 450 to 670 nm, depending on pigment composition (27), whereas plant LHC tend to harvest photons between 420-450 nm and 620-680 nm. Many plants and cyanobacteria also contain small proportions of long-wavelength absorbing Chl *a* molecules (red-chlorophylls) associated with photosystem I (PSI) to extend their light-harvesting capacity in shaded environments (28, 29). However, due to the ancient origins of Chl *a*-based photosynthetic systems, it is difficult to reconstruct the evolutionary history of Chl *a*-based LHC diversification.

By contrast, the comparatively recent origin of Chl *d* tens of millions of years ago in the cyanobacterium *Acaryochloris marina* (30–32) offers the rare opportunity to investigate the process of LHC diversification following the innovation of a novel Chl pigment. Chl *d* differs from Chl *a* by the replacement of a vinyl group with a formyl group at the C-3 position of the chlorin ring, which shifts pigment absorption into the far-red (FR) spectral region (700-750 nm) (30). In *A. marina*, Chl *d* constitutes >90% of Chl content, including the reaction center Chls where photochemistry occurs (33). This enables *A. marina* to thrive in low-light, low-energy habitats, and it is widely distributed in shallow temperate and tropical saline environments where visible light is filtered, often attached to red algae or animals (34). Like prochlorophyte cyanobacteria, *A. marina* and its close relative *A. thomasi*, which does not produce Chl *d* (35), use membrane-bound Pcb proteins for light harvesting (36, 37). Yet, it remains unknown whether the innovation of Chl *d* has been followed by the diversification of the *A. marina* LHC to specialize on different FR wavelengths, as predicted by niche-partitioning theory. Resolving this question has implications for our understanding of the process of biological diversification after the origin of a novel trait.

To investigate LHC functional diversity, we first characterized Chl absorption and fluorescence properties for a collection of *A. marina* laboratory strains for which genome and sequence data are available (34, 38). This revealed distinct *A. marina* spectral phenotypes, which we then mapped to a genome-wide phylogeny to infer the history of *A. marina* LHC diversification. We next considered possible underlying mechanisms of divergence in LHC structure among *A. marina* spectral types. Finally, we investigated how observed phenotypic differences in the *A. marina* photosynthetic apparatus impact photosynthetic oxygen evolution and growth rate in different FR environments. We report that the innovation of Chl *d* was followed by the functional divergence of *A. marina* LHC, with consequences for *Acaryochloris* ecological diversification.

## Results and Discussion

### Divergence of *A. marina* photosynthetic spectral types from an intermediate ancestor

We first obtained room temperature (RT) absorption and fluorescence emission spectra for a sample of 30 *A. marina* laboratory strains isolated from diverse geographic locations and grown in a white light environment (34, 38). The absorption spectra of strains were very similar (Fig. S1), including the characteristic Q_y_ peak of Chl *d* at 710-715 nm, and pigment composition is qualitatively identical among strains with respect to the kinds of chlorophyll and carotenoid pigments produced (Chl *d* with small amounts of Chl *a* and Pheophytin *a*, α-carotene, zeaxanthin; Fig. S2). However, Fig. 1A highlights key differences observed for representative strains. Strain HP10 exhibited an additional shoulder of long-wavelength absorbance at ∼740-750 nm compared with strains MBIC11017 and MU03 (Fig. 1A). This variation in Chl *d* absorption suggests functional differences in light use across *A. marina* strains. Meanwhile, the type-strain MBIC11017, which is the only *A. marina* strain known to produce the light-harvesting pigment phycocyanin (40), additionally produced an absorption peak for this phycobiliprotein at ∼620 nm (Fig. 1A; 39), obscuring the Q_x_ bands of Chl at 605 nm and 650 nm (Fig. 1A).

**Figure 1.**
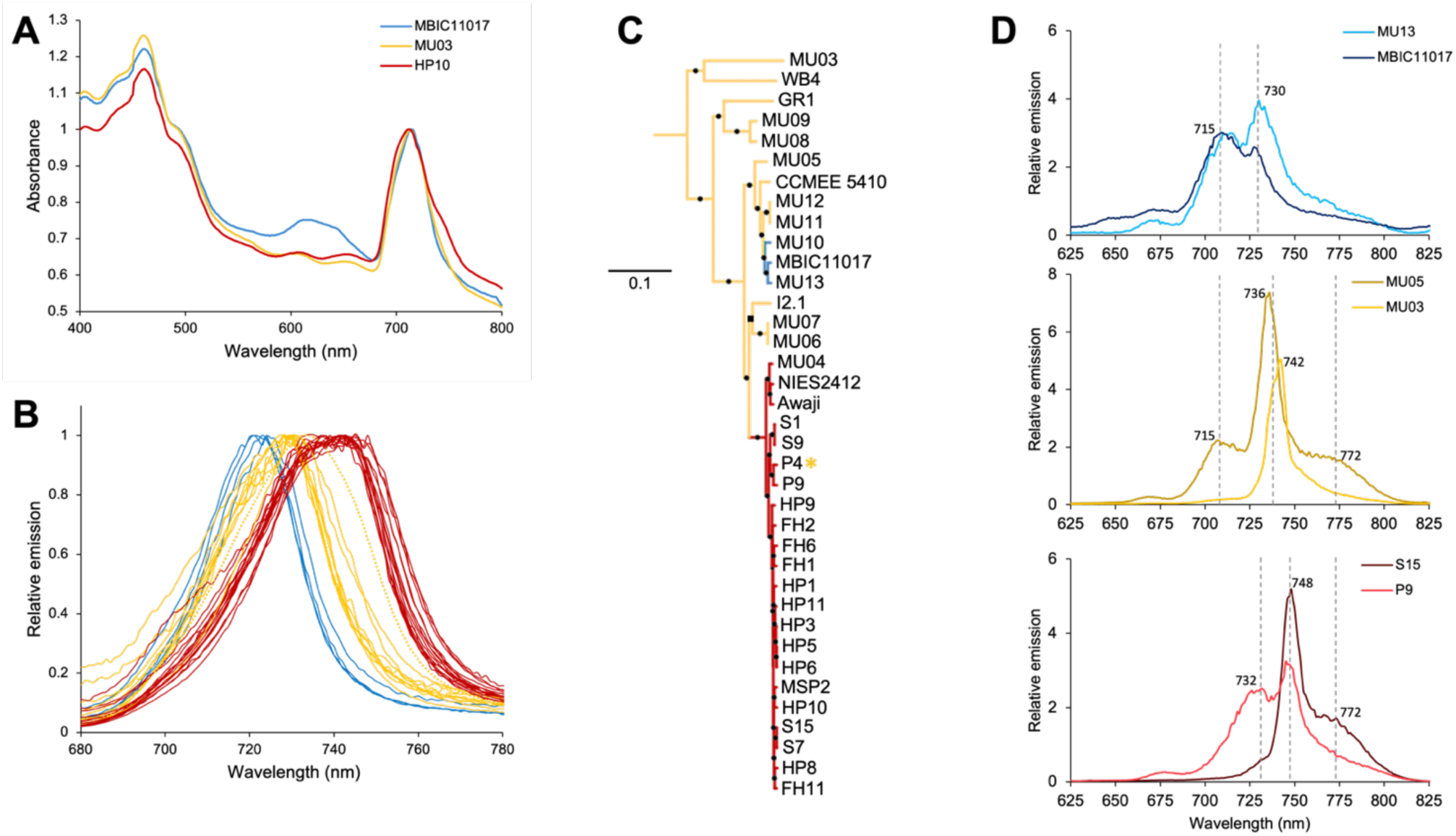
Divergence of *A. marina* photosynthetic spectral types. Data are color-coded by spectral type in all panels. **A**: Absorption spectra of representative strains MBIC11017, MU03, and HP10 normalized by the Chl d peak (696 nm). **B**: RT fluorescence emission spectra of *A. marina* strains grown in white light. P4 emission spectrum indicated by dotted yellow line. **C**: Maximum likelihood phylogeny of *A. marina* strains adapted from (34) that was outgroup-rooted with *A. thomasi* RCC1774. Bootstrap support was >98% for nodes indicated by black squares and 100 % for nodes indicated by black circles. The asterisk by P4 indicates an intermediate phenotype between LW and IW. **D**: 77K fluorescence emission spectra of whole cells from select strains of each spectral type. Dotted lines indicate peaks.

We next characterized RT fluorescence emission spectra, which detect photons radiated by excited electrons of light-harvesting antenna Chl molecules as they return to ground state (particularly the photosystem II (PSII) antenna when performed at RT; 40). Differences among strains were more apparent for RT fluorescence emission compared with absorption (Fig. 1B). We identified three major *A. marina* spectral types: a previously unidentified intermediate wavelength (IW) type with an emission maximum at 727-731 nm; a short-wavelength (SW) spectral type with an emission maximum at 721-724 nm, as reported for *A. marina* strain MBIC11017 (39); and a long-wavelength (LW) spectral type with a broader emission peak and maximum at 738-748 nm, first observed for *A. marina* strain Awaji (41).

Many photosynthetic organisms have the capacity to adjust the composition of their light-harvesting systems to better match the quality of light in the environment by a process called complementary chromatic acclimation (CCA; 42). Several CCA mechanisms have been characterized in cyanobacteria (43, 44). To address whether differences in CCA could explain the different *A. marina* spectral types, we grew strains at different light intensities in both white and FR light. Although strains did exhibit minor shifts in fluorescence emission maximum in response to changes in light quantity and quality, strains consistently remained in the same spectral type grouping irrespective of light environment (Fig. S3). We conclude that the *A. marina* spectral types are due to fixed, evolved differences among strains rather than the product of differences in the regulation of a variable acclimation response.

To understand the evolutionary origins of the divergent spectral types, we mapped the RT fluorescence emission phenotype onto a genome-wide *A. marina* phylogeny (Fig. 1C; 34). Members of the earliest branching lineages exhibited the IW spectral type, indicating that this was most likely the ancestral phenotype. Ancestral state reconstruction (ASR) by the marginal posterior probabilities approximation (MPPA) maximum likelihood model and F81 model for character evolution strongly supports this hypothesis (99.4% marginal posterior probability). Strikingly, the origins of both SW and LW spectral types appear to be comparatively recent (Fig 1C). The SW spectral type (observed in strains MU13, MBIC11017, and MU10) originated within a clade of primarily tunicate-associated *A. marina* that also includes MU11 and MU12 (Fig. 1C; 34), whereas the LW spectral type was innovated by the ancestor of a clade of strains isolated from diverse locations, including western North America, Portugal, Italy and Japan. One strain (P4) exhibits an intermediate phenotype between LW and IW spectral types, suggesting a partial reversion to the ancestral phenotype (Fig. 1B,C; Fig S5A).

### Possible sources of functional variation among *A. marina* spectral types

The observed differences in RT fluorescence emission maxima among spectral types suggest the presence of different energy forms of Chl in the light-harvesting apparatus (40), which impacts the wavelengths of photons that are harvested as well as the flow of excitation energy to the reaction centers (45, 46). To investigate this in more detail, we obtained 77K fluorescence emission spectra, which better resolve functional differences among photosynthetic systems by eliminating biochemical and physiological processes, slowing down energy transfer, and sharpening spectral peaks through the loss of intramolecular vibrations (40). 77K spectra corroborated RT observations of differences among spectral types in Chl energy levels but also revealed variation within each spectral type that were not apparent for PSII-dominated RT fluorescence (Fig. 1D). Both SW and IW spectral types contained a 715 nm and a prominent ∼730 nm emission peak; however, the latter peak was shifted to longer wavelengths in IW strains (ranging from 736-742 nm) (Fig. 1D). MU05 had an additional emission shoulder at 772 nm as well as the 715 nm peak, both of which MU03 (earliest branching *A. marina* strain) lacked, indicating variation within spectral types that was not observed at RT (Fig. 1D). This was also the case for LW strains, yet both S15 and P4 contained a longer-wavelength shift of the prominent peak to 748 nm (Fig. 1D).

Although a detailed characterization of the mechanisms that give rise to this rich functional variation within and between *A. marina* spectral types is the subject of future work, we identify possible contributors to these patterns, which are not mutually exclusive. The different spectral types use the same kinds of Chls and carotenoids (Fig. S2); therefore, differences among spectral types are likely due to (i) changes in protein environment (e.g., hydrophobicity) in the vicinity of Chl molecules that are caused by variability in Pcb sequence and/or composition within light-harvesting complexes; and/or (ii) differences in the amounts of Chl *d* and Chl *a* molecules in light-harvesting antenna proteins themselves.

*A. marina pcb* genes are homologs of prochlorophyte *pcb* and are members of a large gene family that includes the iron-stress gene *isiA* and the *psbC* and *psbB* genes of PSII (Fig. S4; 47, 48). Whereas *A. marina pcbA* genes are more closely related to *isiA*, *pcbC* paralogs form a separate clade, and all (*pcbA1*, *pcbA2*, *pcbC*) are derived from three ancestral copies that are found in *A. thomasi* RCC1774 (Fig. 2; Fig. S4), most likely obtained by ancient horizontal transfer from prochlorophyte(s). *A. marina* strains exhibit extensive variation in *pcb* copy number (ranging from 6 to 10 copies; Fig. 2A) due to a dynamic history of gene duplication, differential retention of paralogs and horizontal acquisition of genes from other *A. marina* strains (Fig. 2). Basal *A. marina* lineages diverged prior to several *pcb* duplication events and consequently harbor fewer gene copies (Fig. 2A, B), which is associated with fewer fluorescence emission peaks (Fig. 1D).

**Figure 2.**
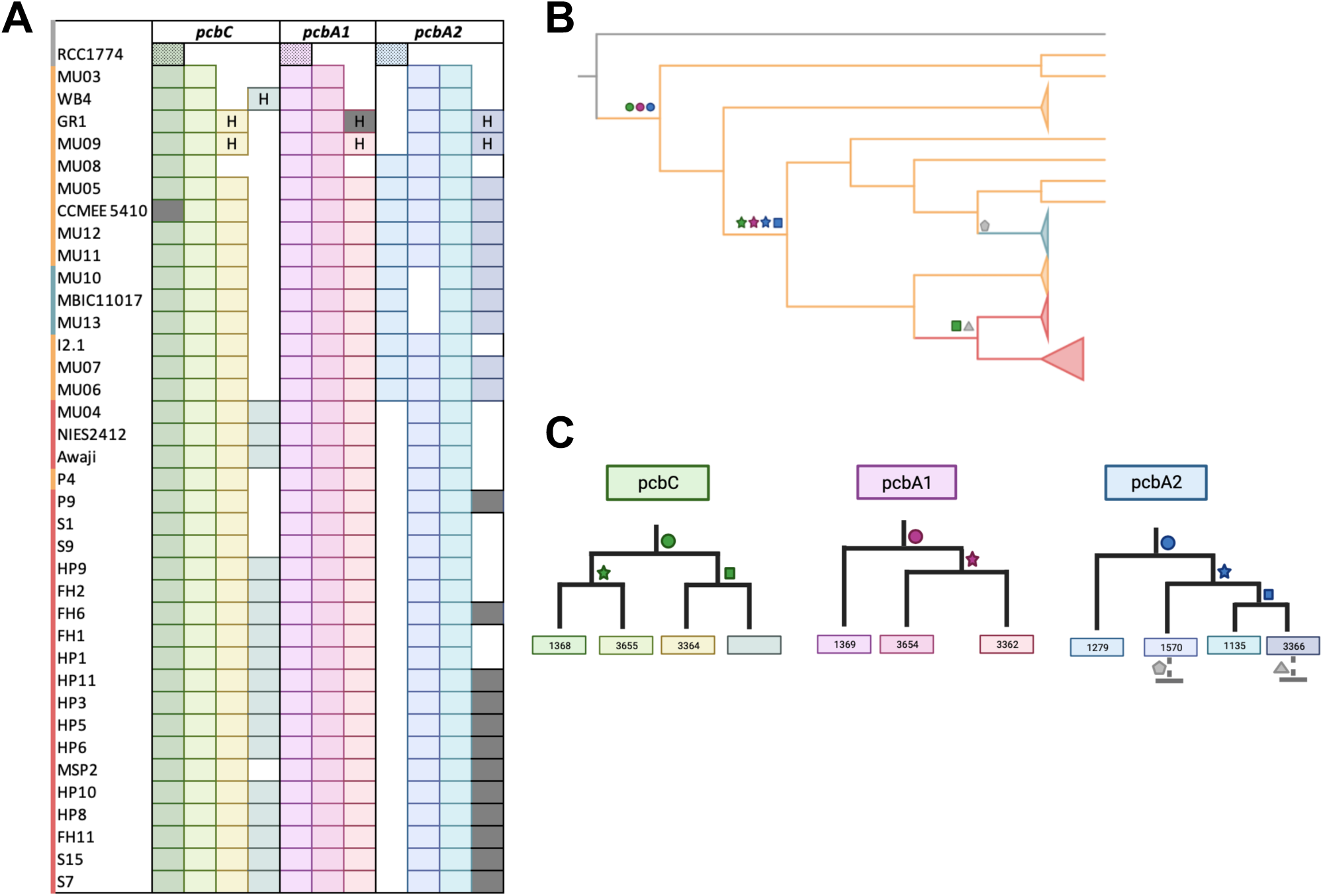
Pcb copy number variation and complex evolution in *Acaryochloris.* **A**: Presence/absence table of *pcb* gene copies in *Acaryochloris* strains. Strains are color-coded by spectral type and are ordered as in Fig. 1C. Gray boxes indicate pseudogenes and white boxes indicate the absence of a gene copy. Horizontally acquired genes are indicated by “H”. **B**: Partially collapsed *Acaryochloris* phylogeny outgroup-rooted with *A. thomasi* RCC1774 (gray). *A. marina* branches are colored by identified spectral type. Symbols indicate inferred duplication events or losses. **C**: Inferred gene trees of *pcbC*, *pcbA1*, and *pcbA2* copies. Green, pink, and blue symbols indicate a duplication, gray symbols indicate a loss as in **B**.

Although the same *pcb* repertoire is generally shared within each spectral type (Fig. 2A), recombination of *pcb* alleles during *A. marina* diversification may contribute to spectral differences between closely related strains. For example, strains P4 and P9 possess the same number and kinds of Pcbs (Fig. 2A), but P4 alleles for two *pcbA* loci are more closely related to those of basal strains than LW strains (Fig. S5). This may contribute to the unique RT fluorescence emission spectrum of strain P4, which is intermediate between ancestral and LW spectral types (Fig. 1B; Fig. S5).

Differences in Pcb stoichiometry may also contribute to functional differences of *A. marina* LHC. We purified Chl-protein complexes of selected strains by sucrose density gradient centrifugation (Fig. 3A) and characterized fluorescence properties of specific fractions. Fraction F3 primarily contained PSI complexes based on position in the density gradient and similarity of pigment absorption and emission compared with published results for iron-replete MBIC11017 (37, 49, 50). Notably, the F3 absorption spectrum for SW strain MBIC11017 is missing a prominent longer-wavelength shoulder that is observed in MU05 and P4 (Fig. 3B); this FR shoulder was further resolved at 77K as a broad band of intense fluorescence emission at wavelengths greater than 750 nm, which was absent in MBIC11017 (Fig. 3C). This broad bandwidth suggests the presence of multiple Chl molecules with lower energy than the PSI reaction center (peak wavelength of 740 nm = 1.68 eV; 51)), which would necessitate thermally activated uphill energy transfer from the antenna to the PSI special pair. Low energy Chls (red-chlorophylls) within other cyanobacterial PSI antenna have been shown to be involved in light harvesting (reviewed in 52). Liquid chromatography-mass spectrometry (LC-MS) analysis of F3 for MBIC11017 and MU05 showed enrichment of PSI proteins together with multiple Pcb proteins. The Pcbs of MBIC11017 (SW) were dominated by PcbA1 paralogs 1369 and 3654 (MBIC11017 numbering; Table S1; Fig. 2C). PcbA1 3654 expressed in Chl *a*-producing *Synechocystis* sp. PCC 6803 exhibits a 723-nm emission peak at 77K (53); we therefore assigned the MBIC11017 emission band at 729 nm (red-shifted due to the presence of Chl *d*) to these PcbA1 copies. By contrast, F3 of MU05 contained similar amounts of PcbC copies 1368/3655 (these paralogs have identical sequences) compared with 1369 and 3654 (Table S2). These Pcbs may therefore be responsible for the long-wavelength fluorescence that is absent in MBIC11017, and the transition from IW to SW spectral types most likely involved evolved differences in Pcb stoichiometry of the PSI light-harvesting complex.

**Figure 3.**
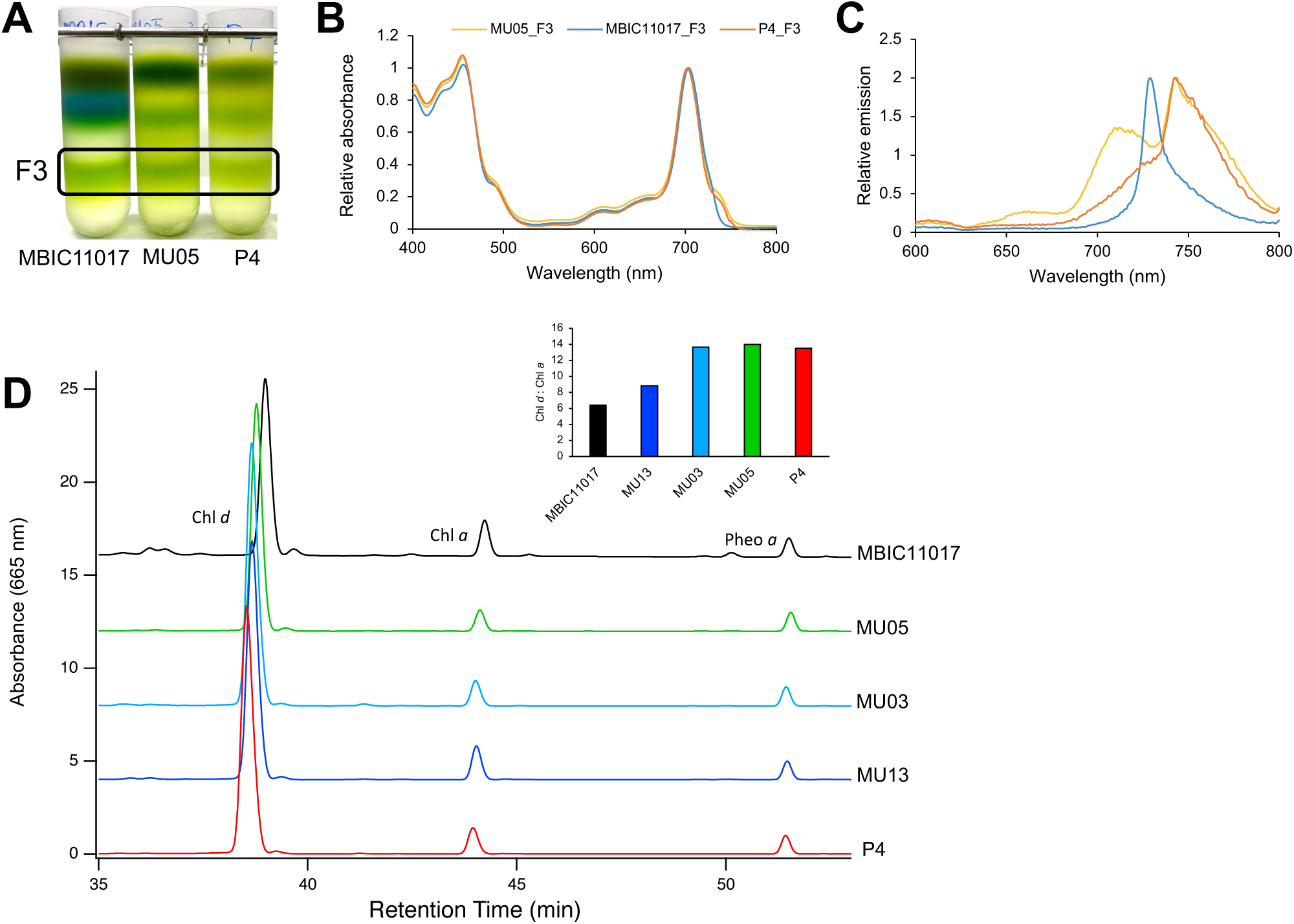
Possible contributors to derived shifts in *A. marina* spectral types. **A:** Fractions of Chl-protein complexes from MBIC11017, MU05, and P4 after sucrose density gradient centrifugation. Fraction 3 (F3) was further purified, and absorption (**B**) and 77K fluorescence emission (**C**) spectra were performed for F3 fractions of MU05 (yellow), MBIC11017 (blue), and P4 (orange). **D**: HPLC chromatogram of *A. marina* strains MBIC11017, MU05, MU03, MU13, and P4. Inset: Estimated ratios of Chl *d*: Chl *a*, when normalized to Pheophytin *a*.

Finally, differences in Chl *d*: Chl *a* in light-harvesting antenna proteins may contribute to the transition from IW to SW spectral types. Pheophytin *a*-normalized Chl *d*: Chl *a* of IW and LW strains (13.7 ± 0.14) compared to SW (7.61 ± 1.23) indicated relatively greater antenna Chl *a* in SW strains (Fig. 3D). Pheophytin *a* is the primary electron acceptor of PSII and should not vary across strains (54). Higher Chl *d*: Chl *a* in IW and LW strains potentially contribute to lower energy traps, and the transition from IW to SW most likely involved increased antenna Chl *a* molecules.

### Physiological and fitness consequences of spectral type divergence

Do differences in spectral type have phenotypic consequences, such as specialization on different FR wavelengths? Following the evolution of Chl *d* and the ability to access new FR wavelengths, we predicted that a basal *A. marina* strain with ancestral spectral type would exhibit a generalist strategy. During the process of diversification as a response to variable selective pressures, new ecological opportunity, or competition avoidance with other *A. marina* strains, they might subsequently partition ecological niche space through spectral type divergence and specialization on specific FR wavelengths (55, 56). To investigate this, we compared growth rates and photosynthetic oxygen production of representative strains from each spectral type in different FR environments provided by LED sources with peak emission at 706 nm (approximately the Chl *d in vivo* absorption maximum) and 731 nm, respectively).

Strains responded to the light environments differently (Fig. 4A), as indicated by a highly significant strain x light interaction term in a linear model (*F*_2,15_ = 12.43; *P* = 0.001). Specifically, basal IW strain MU03 exhibited no difference in growth rate between light environments (*F*_1,4_ = 0.30; *P* = 0.62; Fig. 4A), as expected for an intermediate generalist phenotype. By contrast, derived spectral types exhibited growth patterns consistent with predictions based on evolved shifts in Chl fluorescence (Fig. 4A): HP10 (LW) grew significantly slower in 706 nm compared to 731 nm (*F*_1,4_ = 17.2; *P* = 0.01), whereas the growth rate for MU13 (SW) was faster in 706 nm (*F*_1,4_ = 6.02; *P* = 0.07). Observed differences in absolute performance among strains may reflect phenotypic differences with respect to their responses to other environmental parameters in the growth assay (e.g., temperature, light intensity). With respect to photosynthetic oxygen evolution, LW strain HP10 exhibited a much lower rate at 706 nm (Fig. 4B; *F*_2,27_ = 4.91, *P* = 0.016 for the strain x light interaction term) than either MU03 (IW) and MU13 (SW). The low net rates of oxygen evolution for HP10 in 706 nm light might explain, in part, the reduced growth rate we observed at shorter wavelengths.

**Figure 4.**
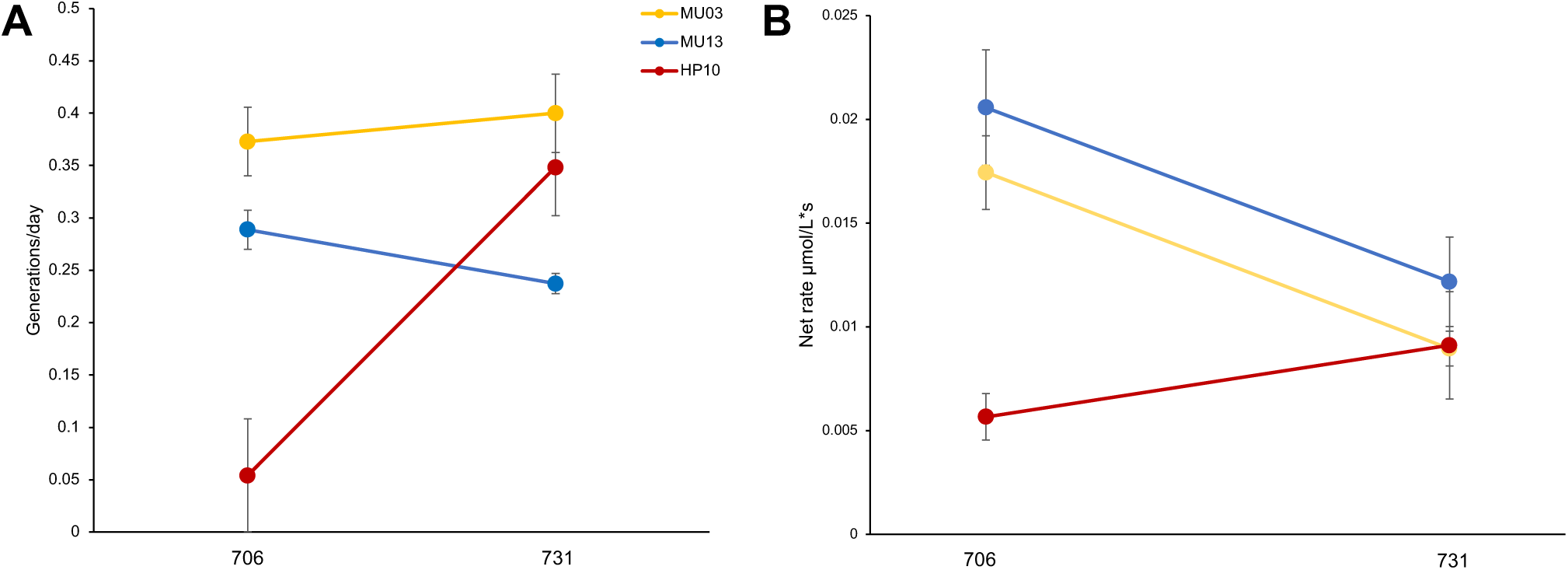
Physiological and fitness consequences of spectral type divergence. Error bars are standard errors **A**: Population growth rates in 706 nm and 731 nm FR light environments for MU03 (IW, yellow), MU13 (SW, blue), and HP10 (LW, red). **B**: Net rates of oxygen production for MU03, MU13, and HP10 when grown and assayed in 706 nm and 731 nm, respectively.

Taken together, *A. marina* spectral types exhibit physiological differences when grown in different FR light environments, suggesting that derived spectral types have specialized to use distinct FR wavelength environments. Divergence of *A. marina* spectral types therefore resembles the diversification of beak size and shape in Darwin’s finches that is associated with specialization on different diets (57). *A. marina* strains of different spectral types are known to co-occur (e.g., P4/P9 and MU03/MU04; 34), suggesting that niche partitioning via character displacement may have been favored by competition avoidance. We conclude that, following the innovation of a Chl *d*-based light harvesting system in *A. marina*, resource partitioning has enabled strains to specialize to use different FR wavelengths for photosynthesis.

## Methods

### *Acaryochloris* culture maintenance

*Acaryochloris* strains were cultured in IO BG-11 or modified IO BG-11 media as in (38). Batch cultures containing 75 mL of media were maintained in 250 mL Erlenmeyer flasks at 20 °C or 30 °C under 12 h cycles of 20-25 μmol photons m^−2^s^−1^ cool white fluorescent light (350 – 800 nm). All strains are maintained in the University of Montana Culture Collection for Cyanobacteria and are available upon request.

### Pigment extraction, room temperature spectroscopy, and tube assays

Absorbance scans were performed *in vivo* by first normalizing by optical density at 750 nm (OD_750_) and measuring absorption from 300-800 nm with a Beckman Coulter DU 530 spectrophotometer (Indianapolis, IN). To calculate Chl *d* concentration, a methanol assay was performed as in (38, 58). Briefly, 2 mL of culture were harvested and pelleted by centrifugation. The supernatant was discarded, and cell pellets were resuspended in 2 mL ice cold 100 % methanol. To extract pigments, samples were stored on ice in the dark for ∼30 min, after which they were centrifuged to pellet cell debris. Chl *d* concentration (μg/ml) was determined by measuring the absorbance at 697 nm and then using the published extinction coefficient of Chl *d* in methanol (63.68 x 10^3^ L mol^−1^cm^−1^; 59). Aliquots of each culture were normalized by Chl *d* concentration before taking fluorescence emission spectra (450 nm excitation, 475-800 nm scan) with a Photon Technology International model QM-7/2005 spectrofluorometer (Ontario, Canada). Spectra were further normalized by peak wavelength.

Select *A. marina* strains were aliquoted into 15 mL of IOBG-11 media in culture tubes with a starting OD_750_ of 0.01. Tubes were propagated into different intensities of FR (0.94, 1.7, 3.5, and 10 μmol photons m^−2^ s^−1^) and cool white light (15, 30, 46, 70, and 99 μmol photons m^−2^ s^−1^) at 20 °C. Metal screening was used to reduce light intensity. Pigments were extracted to normalize to Chl *d* concentration before fluorescence emission spectra were collected as above.

### Phylogenetic analysis of *A. marina* LHC

The *Acaryochloris* spp. maximum likelihood phylogeny was used from (34), with *Cyanothece* sp. PCC7425 (NCBI accession GCA_000022045.1) as the outgroup. Ancestral state reconstruction was performed with PastML (60) using the marginal posterior probabilities approximation (MPPA) maximum likelihood model and F81 model for character evolution (61). Amino acid sequences of *A. marina* Pcb obtained from annotated genome sequence assemblies in addition to other publicly available cyanobacterial Pcb, CP43, CP47, and IsiA protein sequences (NCBI accessions: PRJNA224116, PRJNA13548, PRJNA158811, PRJNA57995, PRJNA13452, PRJNA213, PRJNA16251, PRJDB11069, PRJNA209725, PRJNA19377, PRJNA246696, PRJEB60184, PRJDB4348, PRJDB5665, PRJNA10645, PRJNA715740, PRJNA28247, PRJNA218025, PRJNA795194, PRJNA12997, PRJNA430151, and PRJNA649288) were used to create a concatenated alignment. We constructed a maximum likelihood tree with 1,000 ultrafast bootstrap replicates (62) using IQ-TREE version 2.0 (63) according to the cpREV+F+I+G4 model of sequence evolution selected by the Akaike information criterion (AIC) in ModelFinder (64). The Shimodaira-Hasegawa-like aLRT (SH-aLRT) was performed in parallel with bootstrap replicates to maximize load balance. To confirm presence/ absence of specific Pcb copies, we manually checked Pcb fragmented sequences for evidence of pseudogenization (premature stop codon) or artifacts from misalignment. The multiple genome alignment tool Mauve (65) was used to examine synteny of operons containing *pcb* copies (Fig. S6). Gene trees of individual *pcb* copies were constructed from nucleotide alignments using the above method according to the TPM2u+F+G4, TIM2+F+G4, TIM2+F+G4 models of sequence evolution for *isiA*, *pcbA1* 1570 and *pcbA2* 1369 copies, respectively.

### HPLC analysis

For HPLC analysis at The Pennsylvania State University, *A. marina* cells were harvested and washed once in 50 mM HEPES/NaOH buffer, pH 7.0 by gentle centrifugation, resuspension, and another centrifugation. Methods were followed as previously described (66, 67). Briefly, pigments were extracted from the cell pellet by sonication in acetone:methanol (7:2, v/v). Protein and other insoluble cell debris were removed by centrifugation, and pigment solutions were filtered through a 0.2-μm polytetrafluoroethylene membrane syringe filter. Next, pigments were analyzed by reversed-phase HPLC on a 25 cm × 4.6 mm analytical Discovery C18 column (Supelco, Bellefonte, PA, USA) using an Agilent Model 1100 HPLC system equipped with a model G1315B diode array detector. Chl ratios were estimated by calculating the area under each curve.

### Membrane separation, fractionation, and spectroscopy

Cell pellets (5-8 g by dry weight) were stored in –80 °C and shipped to The Pennsylvania State University, University Park, for further processing. Pellets were resuspended in MES buffer containing 50mM MES (pH 6.5), 15 mM CaCl_2_, and 10mM MgCl_2_; 10 mL of buffer was used for each gram of cells. After homogenizing cells, a combination of approaches was used for cell lysis. Cells were passaged three times through a chilled French press (pressure ∼ 138 MPa), sonicated, and passed through a microfluidizer (Microfluidics M-110EH-30 Microfluidizer Processor) at 30,000 psi. Further, a volume of 1 mL was mixed with an equal volume of 0.1-mm glass beads and cells were broken by eight rounds of 30 s of bead beating in a Mini-BeadBeater (BioSpec Products). Samples were centrifuged at 4000 rpm for 15 min in 10 °C to remove cell debris. The supernatant was centrifuged at 40,000 rpm for 1 h in 4 °C and pelleted membranes were resuspended in MES buffer. The Chl concentration was measured and adjusted to 0.4 mg Chl ml^−1^ by dilution with MES buffer. Membranes were then solubilized by addition of *n*-dodecyl-β-d-maltoside detergent (DM) to a final concentration of 1% (w/v) and incubated at 4 °C for 1 h. Solubilized membranes were separated from insoluble debris by centrifugation at 8000 rpm for 15 min in 4 °C. The solubilized membranes were loaded onto 5 to 20% (w/v) sucrose gradients containing 0.1% DM in MES buffer, and the gradients were centrifuged for about 18 hours at 108,000*g* in 4 °C. Green-colored fractions were collected from sucrose gradients (see Fig. 3A), dialyzed against MES buffer and concentrated using Millipore Centriprep 100K Centrifugal Filtration Devices (EMD Millipore, Darmstadt, Germany). For further purification, F2 and F3 fractions were loaded onto 5 to 20% (w/v) sucrose gradients containing 0.1% DM in MES buffer and subjected to a second ultracentrifugation for 18 hours at 108,000*g* in 4 °C. Fractions were collected and dialyzed against MES buffer.

Absorption and low-temperature (77K) fluorescence spectra were collected for membrane fractions after normalizing for Chl concentration. Absorption spectra (300-800 nm) were measured with Cary 14 spectrophotometer that was modified for solid-state operation by On-line Instrument Systems Inc., (Bogart, GA, USA). 77K fluorescence emission spectra were measured using an SLM Model 8000C spectrofluorometer that has been modified for computerized solid-state operation by On-line Instrument Systems Inc., (Bogart, GA, USA). Membrane fractions were diluted in MES buffer to normalize Chl concentration and then glycerol was added to produce a final concentration of 60% (v/v). After loading into the measuring tube, samples were quickly frozen in liquid nitrogen. To measure the fluorescence emission from Chl-protein complexes, samples were excited at 450 nm and emission was measured over the range of 600-800 nm.

### LC-MS Preparation

To precipitate the proteins from purified fractions, 12% PEG (w/v) was added to each tube, inverted to mix, and stored in 4 °C overnight. The PEG/protein mixtures were centrifuged for 20 min at 30,000 rpm in 4 °C. Supernatants were decanted, and pellets were briefly dried. The protein pellets were resuspended in resuspension buffer composed of 50 nM MES pH 6.5, 10 mM MgCl_2_, 15 mM CaCl_2_, 10 % glycerol, and 0.05% DM. Samples were stored in –80 °C and prepared for LC-MS performed at the Penn State Proteomics and Mass Spectrometry Core Facility, University Park, PA.

### FR growth and net oxygen production experiments

For each strain, triplicate independent cultures derived from the same inoculum were grown in 706 nm and 731 nm LED boxes in 25 °C. The intensity of LED lights varied between 1.9-2.1 μmol photons m^−2^s^−1^. Growth was measured every 48 hours by taking OD_750_ readings of a 2 mL subsample, after which cultures were randomly moved to different positions within each respective light environment to mitigate any differences in light exposure. Growth rates (generations/day) were estimated from the exponential growth phase of each culture.

Aliquots of exponentially growing cells were gently homogenized and pelleted via low-grade centrifugation (5500*g* for 30 min) and resuspended in 2 mL IO BG-11 media to a normalized Chl *d* concentration. The sample was kept in the dark for at least 5 min prior to measuring net oxygen evolution with a Unisense OX-Eddy Clark-type oxygen sensor when the sample was exposed to 706 nm or 731 nm LED light for 30 sec. Each sample was measured twice and was dark acclimated as well as carefully homogenized with a sterile needle in between measurements. The oxygen sensor signal was recorded every second using the Unisense SensorSuite Logger software. Prior to experimental procedures, the oxygen sensor was calibrated following manufacturer’s instructions for conversion from raw sensor signal (mV) to oxygen concentration (µmol O_2_ L^−1^). For each of the light treatments, the rate of oxygen evolution (µmol O_2_ L^−1^ s^−1^) was estimated by the initial slope of a stable increase of oxygen over 10-15 seconds.

### Quantification and statistical analysis

Differences in growth rates and net rates of oxygen evolution of *A. marina* strains MU03, MU13, and HP10 in both light environments were assessed by linear models in JMP version 17.2 (SAS Institute Inc., Cary, NC). Details of other quantification analyses can be found in the Results and Discussion text.

