## Supplementary Information for "Niche partitioning in a cyanobacterium through divergence of its novel chlorophyll *d*-based light-harvesting system"

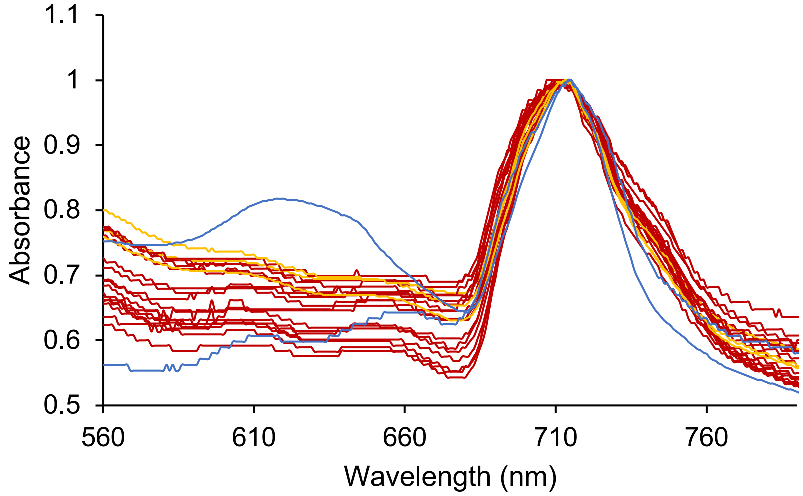


**Figure S1**. Absorbance spectra of a subsample of *A.marina* strains in white light normalized by the Chl d peak (696 nm). Strains are colored by spectral type: LW (red), IW (yellow), and SW (blue).

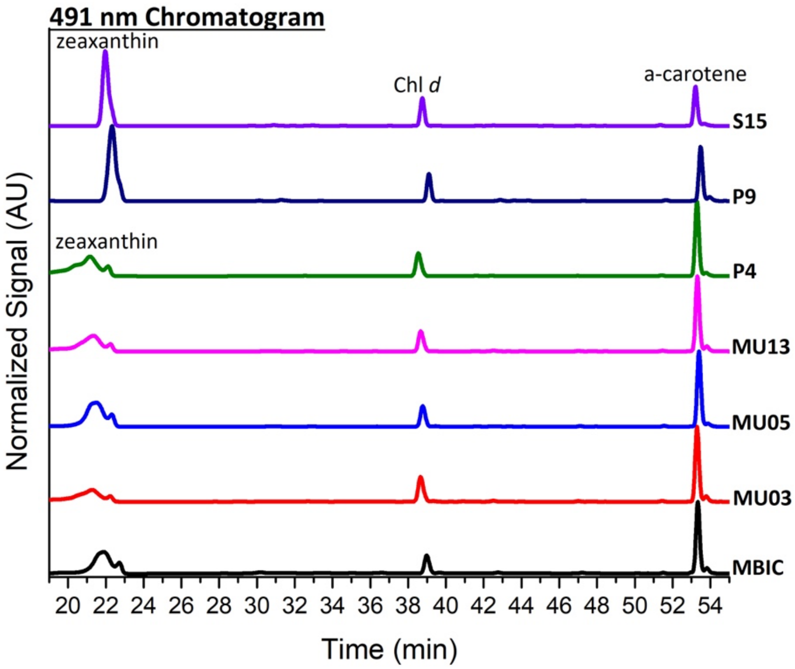


**Figure S2**. HPLC chromatogram of select *A. marina* strains, showing relative signals of zeaxanthin, α-carotene, and Chl *d*.

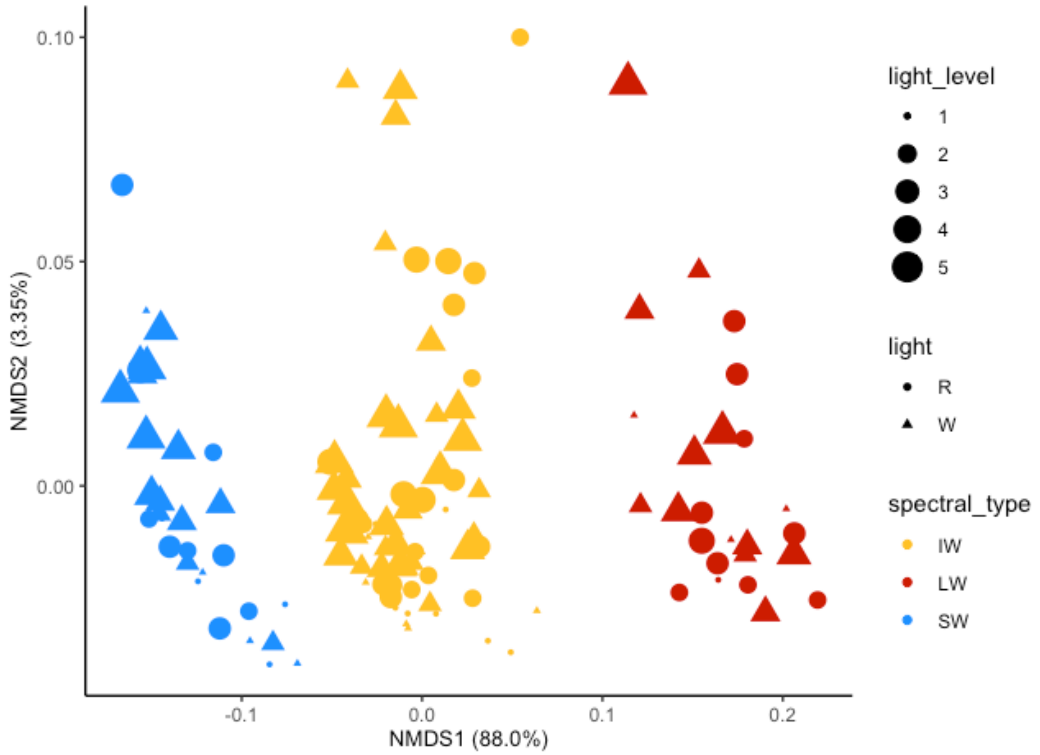


**Figure S3**. NMDS of RT fluorescence emission spectra from *A. marina* strains grown different light quantity and quality. Strains were grown in different levels of FR and white light, signified by triangles and circles, respectively. Strains are colored by their spectral type: LW (red), IW (yellow), and SW (blue). Bray-Curtis dissimilarity indexes were calculated from normalized fluorescence emission spectra and plotted as a non-metric multidimensional scaling (NMDS) ordination in R.

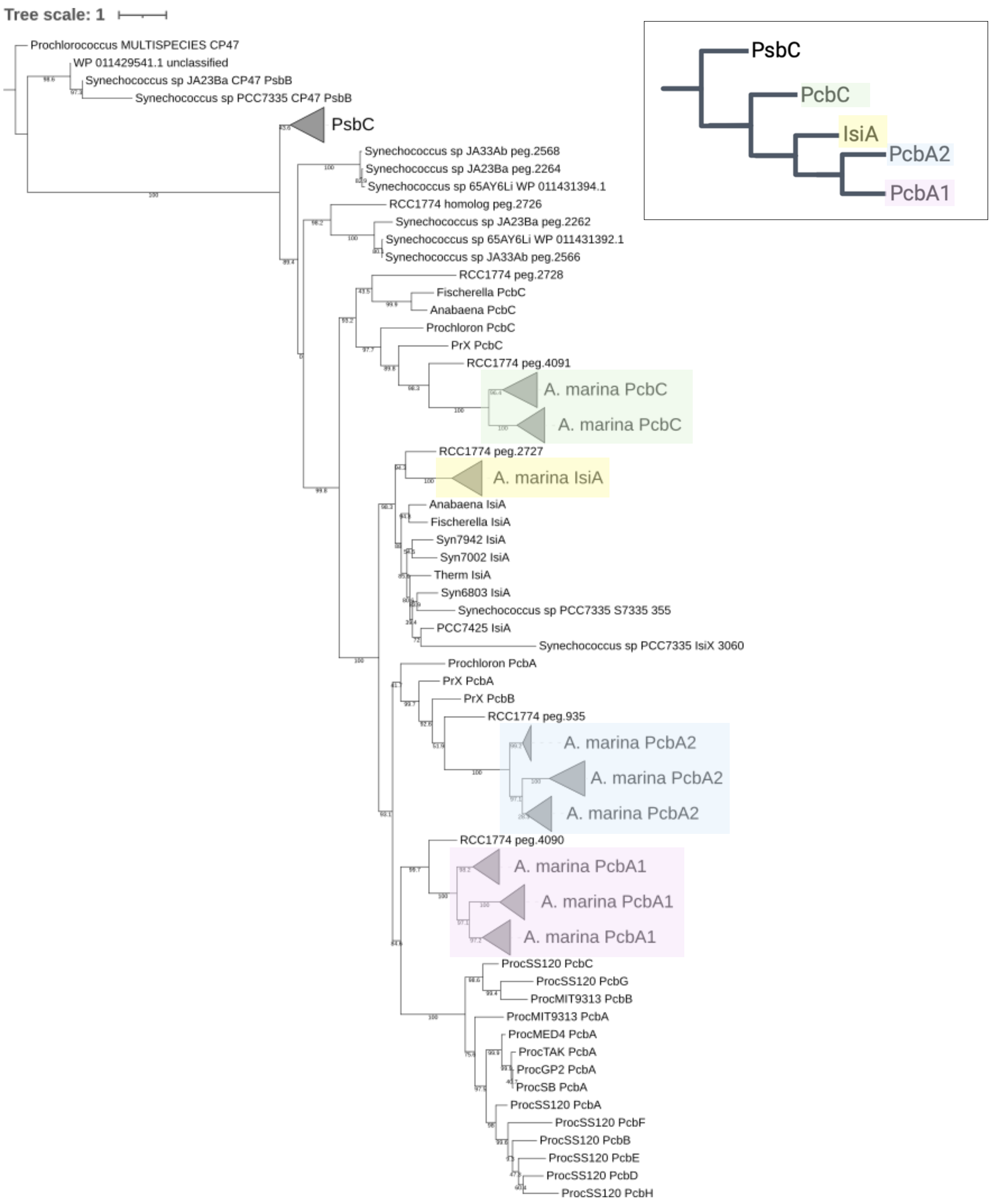


**Figure S4**. Maximum likelihood phylogenetic reconstruction of Pcb proteins. Bootstrap (n=1000) support is shown for each node. *A. marina* Pcbs are indicated by colored boxes. Tree scale indicates substitutions per site.

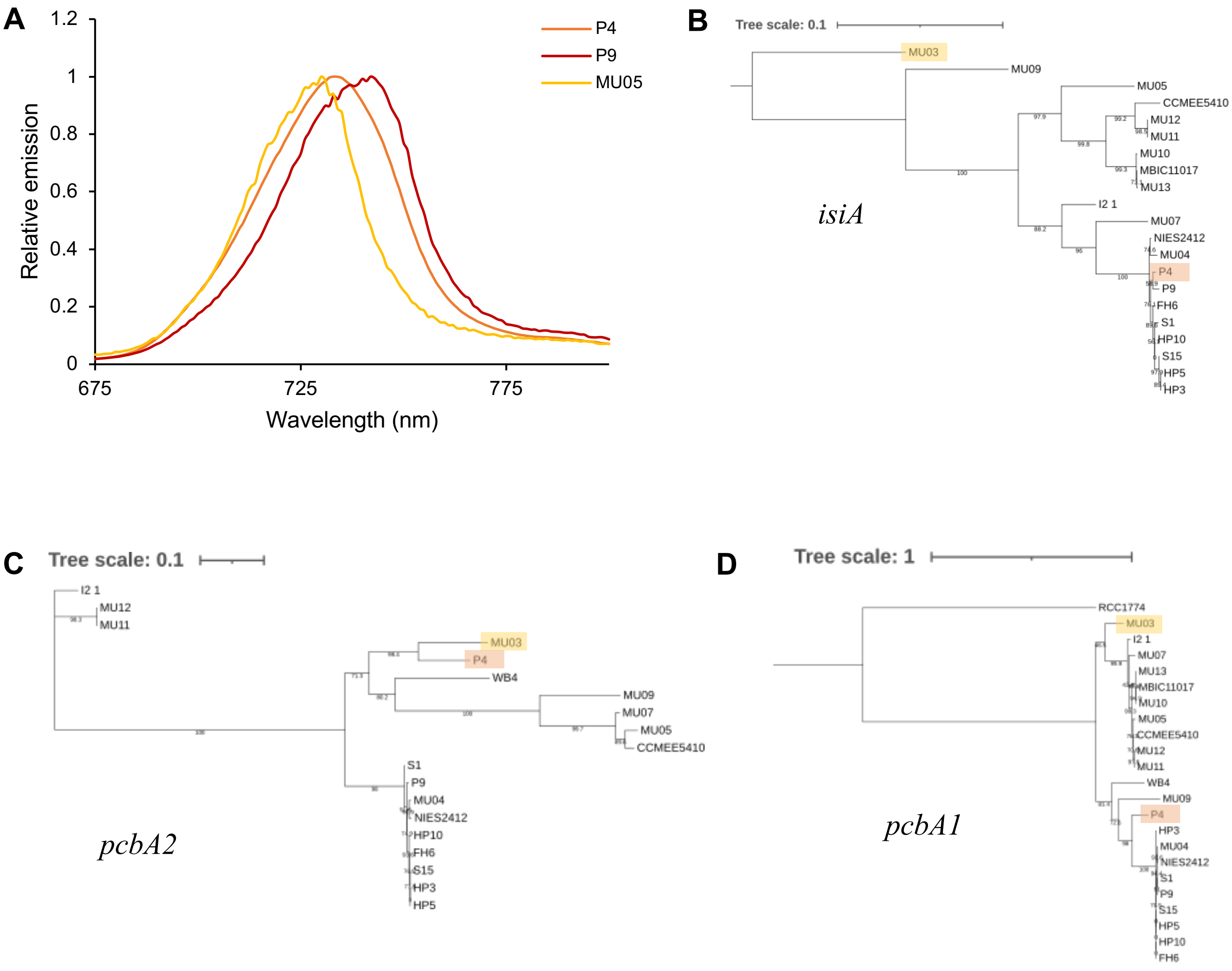


**Figure S5**. **A**: RT fluorescence emission spectra of P9 (LW, red), MU05 (IW, yellow) and P4 (orange). P4 exhibits a unique intermediate fluorescence emission spectrum that is between LW and IW spectral types. Maximum likelihood phylogenetic reconstructions of genes *isiA* (**B**), *pcbA2* 1570 copy (**C**) and *pcbA1* 1369 copy (**D**). Bootstrap (n=1000) support is shown for each node. Whereas the *isiA* gene tree reflects the *A. marina* species tree, the *pcbA2* and *pcbA1* copies have different topology, in which P4 does not group with P9. The earliest branching *A. marina* strain MU03 is highlighted in yellow and P4 is highlighted in orange.

**Table S1**. LC-MS: Select proteins identified in F3 fraction in strain MBIC11017 (SW)

| **Protein name** | **Accession** | **Coverage (%)** | **Theoretical MW (kDa)** | **Unique peptide** | **Sequest Score** |
|---|---|---|---|---|---|
| photosystem I core protein PsaA | ABW27465.1 | 19 | 83.3 | 13 | 71.66 |
| photosystem I core protein PsaB | ABW27466.1 | 20 | 82.1 | 15 | 79.51 |
| photosystem I protein PsaL | ABW26465.1 | 23 | 15.3 | 4 | 15.99 |
| photosystem I protein PsaK | ABW26658.1 | 45 | 9.3 | 4 | 14.94 |
| **Chlorophyll *a*/*b* binding protein PcbC (3655/1368)** | **ABW28645.1** | **8** | **37.9** | **2** | **7.11** |
| **Chlorophyll *a*/*b* binding protein PcbA1 (1369)** | **ABW26401.1** | **14** | **38.7** | **6** | **19.61** |
| **Chlorophyll *a*/*b* binding protein PcbA1 (3654)** | **ABW28644.1** | **15** | **38.7** | **6** | **19.5** |
| photosystem II protein PsbK | ABW28836.1 | 31 | 4.9 | 3 | 8.34 |
| photosystem II protein D1 PsbA | ABW27180.1 | 8 | 39.6 | 1 | 10.68 |
| photosystem II CP47 protein PsbB | ABW27041.1 | 7 | 56.4 | 3 | 16.63 |

**Table S2**. LC-MS: Select proteins identified in F3 fraction in strain MU05 (IW)

| **Protein name** | **Accession** | **Coverage (%)** | **Theoretical MW (kDa)** | **Unique peptide** | **Sequest Score** |
|---|---|---|---|---|---|
| photosystem I core protein PsaA | peg.2727 | 21 | 83.4 | 14 | 81.13 |
| photosystem I core protein PsaB | peg.2726 | 8 | 82 | 9 | 53.61 |
| photosystem I protein PsaL | peg.347 | 16 | 15.4 | 4 | 13.79 |
| photosystem I protein PsaK | -- | -- | -- | -- | -- |
| **Chlorophyll *a*/*b* binding protein PcbC (3655/1368)** | **peg.2043** | **18** | **38.6** | **5** | **22.59** |
| **Chlorophyll *a*/*b* binding protein PcbA1 (1369)** | **peg.320** | **12** | **38.7** | **4** | **14.94** |
| **Chlorophyll a/*b* binding protein PcbA1 (3654)** | **peg.804** | **11** | **38.7** | **3** | **12.35** |
| photosystem II protein PsbK | peg.1395 | 31 | 4.9 | 3 | 8.7 |
| photosystem II protein D1 PsbA | peg.4283 | 6 | 40 | 1 | 7.51 |
| photosystem II CP47 protein PsbB | -- | -- | -- | -- | -- |
